# A CATH domain functional family based approach to identify putative cancer driver genes and driver mutations

**DOI:** 10.1101/399014

**Authors:** Paul Ashford, Camilla S.M. Pang, Aurelio A. Moya-García, Tolulope Adeyelu, Christine A. Orengo

**Affiliations:** University College London, Institute of Structural and Molecular Biology, London, UK; Laboratorio de Biología Molecular del Cáncer, Centro de Investigaciones Médico-Sanitarias (CIMES), Universidad de Málaga, Málaga, Spain

## Abstract

Tumour sequencing identifies highly recurrent point mutations in cancer driver genes, but rare functional mutations are hard to distinguish from large numbers of passengers. We developed a novel computational platform applying a multi-modal approach to filter out passengers and more robustly identify putative driver genes. The primary filter identifies enrichment of cancer mutations in CATH functional families (CATH-FunFams) – structurally and functionally coherent sets of evolutionary related domains. Using structural representatives from CATH-FunFams, we subsequently seek enrichment of mutations in 3D and show that these mutation clusters have a very significant tendency to lie close to known functional sites or conserved sites predicted using CATH-FunFams. Our third filter identifies enrichment of putative driver genes in functionally coherent protein network modules confirmed by literature analysis to be cancer associated.

Our approach is complementary to other domain enrichment approaches exploiting Pfam families, but benefits from more functionally coherent groupings of domains. Using a set of mutations from 22 cancers we detect 151 putative cancer drivers, of which 79 are not listed in cancer resources and include recently validated cancer genes EPHA7, DCC netrin-1 receptor and zinc-finger protein ZNF479.

## Introduction

Advances in technology have made exome and whole-genome sequencing commonplace and have been catalysts for large-scale concerted cancer genome sequencing efforts such as TCGA^1^ and ICGC^2^. In tandem with details of tumour types, histology, treatments and patient outcomes, these sequences provide unique opportunities for identifying which mutations and genes drive tumour expansion and how these vary between cancer types.

Mutations observed in tumours may be drivers, positively influencing tumour progression, or passengers, which are incidental and have no net effect^3^. Methods such as MutSigCV^4^ analyse somatic mutations from tumour samples to identify sequence positions mutated above a significance threshold using a sophisticated model of background mutation rates. However, finding driver genes using individual point mutations lacks the statistical power to uncover many driver genes as the heterogeneous mutation landscape of cancer genomes leads to many genes having few mutations, thus identifying significant point mutations requires many tumour samples^3^. A complementary approach is to analyse mutations by mapping to 3D protein structures: Structural studies can help identify mutations clustering in specific regions of a protein and highlight cases where rare mutations - that may lie far apart in sequence - are close together when mapped to residues in the protein’s structure. Multiple recent methods aid driver gene identification using structure-based algorithms: by calculating frequencies of distances between mutated residue pairs^5^; calculating a clustering coefficient using weighted sums of mutated pairs^6,7^; using permutation testing of mutation distance distributions^8^; finding mutational hotspots within spherical regions^9-12^ and testing protein complexes^13,14^.

Other developments to enhance driver gene detection, that focus on regions in the protein, include clustering of mutations on sequence regions^15^ or using Pfam^16^ protein domains^10,17-19,20^. As evolutionarily-related, discrete & independently folding units of sequence, domains are often found in multiple genes and in different contexts (i.e. multiple domain architectures), therefore domain enrichment may enhance both the statistical power for driver detection and allow clearer prediction of the functional impacts of mutations^21-23^. Sequence hotspots can be detected more easily in enriched domains^10,17,18,24,25^ and can be analysed using co-location with functional sites^24^ such as catalytic sites^26^, phosphosites^27^ and protein-protein interface (PPI) residues^28,29,30-34^. The distribution of cancer mutations to functional sites can be compared with polymorphisms obtained from UniProt^35^. For example, Skolnic *et al*^36^ found an enrichment of disease-causing mutations at functional sites or at ligand-biding sites adjacent to PPIs. Other studies assess how mutation proximity to functional sites varies between known oncogenes and TSGs^6,37^.

Network approaches can also be used to interpret cancer gene sets in terms of cellular processes^38^. In particular, multiple genes containing mutations in a tumour sample may belong to different components of the same biological pathway, and databases such as PathwayCommons^39^ can help predict functional consequences of such mutated gene sets. Previously derived functional interaction networks^40^, built from both known and predicted protein-protein interactions, can be used to analyse driver gene lists for functional consequences, for example, by using GO term enrichment of network modules identified. Additionally, protein-protein interaction networks such as STRING^41^ permit topological predictions, such as whether a particular gene is a hub or how dispersed a set of genes are on the network^42^.

We present a novel study that detects enrichment of cancer mutations in our CATH Functional Families (CATH-FunFams)^43^. CATH v4.0 contains over 25 million domain sequences of known or predicted structure classified into 2,735 homologous superfamilies. Within each CATH superfamily, CATH-FunFams comprise evolutionary related domains^44^ grouped into functionally coherent sets; as such they can help identify functions that appear to be targeted in different diseases. Our work builds on earlier studies that used Pfam domains^45-48^ to detect enrichment, and a recent study that tests enrichment in CATH superfamily domains^49^. Whilst Pfam families provide high-quality annotation of evolutionary relationships, they may also group related proteins whose domains have diverged in function. Recent studies^43,50^ have shown that CATH-FunFams exhibit higher levels of functional purity than either Pfam domains or by using CATH superfamilies. By identifying CATH-FunFams enriched with mutations (‘MutFams’) we filter out mutations that don’t affect protein function (and thus are probably passengers) and as a result identify genes that are more likely to be drivers. In addition, this approach reveals functional rather than domain enrichment.

We show that heatmap clustering of cancers by their MutFams provides sensible groupings and that the top mutated genes from the MutFams appear to map to a specific set of biological processes as they are significantly less dispersed on a protein-protein interaction network compared with sets of randomly chosen genes. Using both GO biological process and functional network enrichment of the top mutated genes from all MutFams we find convergence on cancer-related processes and pathways. In addition, pathway analysis of a comparable set of genes from a related Pfam-based approach^10^ shows that some of the enriched pathway modules are commonly identified by both approaches.

Finally, we performed 3D clustering of all available mutations mapped to representative domain structures of the MutFams, followed by detailed proximity analysis of clusters to various functional sites, on the premise that mutations from different cancers that cluster near the same sites are likely to be having similar functional impacts. We find that, in general, our 3D clustered mutations are closer to functional sites than unfiltered cancer mutations.

Our research thus suggests that finding mutation enriched CATH-FunFam domains is a novel and helpful way to filter out passenger mutations, and that by selection of the most highly mutated genes in each CATH-FunFam we can obtain a broad list of those implicated in cancer. We provide a broad list of 472 genes identified from our MutFams along with a confidence score that can be used to filter for a higher confidence subset of genes. The score reflects other evidence supporting a gene’s involvement in cancer: the predicted functional effects; the identification of cancer-enriched modules on a functional interaction network; significant clustering of mutations on protein structures near functional sites and/or agreement with the Cancer Genome Census (CGC) or Pfam based gene sets. We find 151 genes (in 98 MutFams) are supported by at least one of these confidence tests, with 79 novel genes (in 41 MutFams) not identified, as yet, by the CGC or in the Pfam gene set. The value of our approach is that by filtering mutations to enrich for those with functional effects it could help with identification and prioritisation of driver genes from large-scale cancer genomics studies.

## Results

We analysed somatic, missense mutations from exome/genome-wide studies from 22 cancer types to identify mutationally enriched CATH-FunFam domain families, which we term MutFams. CATH-FunFams are functionally coherent groups of relatives and domains within them share the same patterns of conserved residues unique to the family (i.e. functional determinants such as ligand binding and specificity determining residues)^43^. This step identifies mutationally enriched domain families associated with a more specific function than by testing for enrichment at the domain superfamily level (either with CATH or Pfam). Each CATH-FunFam has at least one experimentally characterised relative with GO functional annotation.

We compared mutated genes obtained from MutFams with a subset of cancer genes from the Wellcome-Sanger Cancer Gene Census (CGC)^51^ containing somatic missense mutations known to be associated with cancer. We also compared with another comprehensive set of genes identified using a related Pfam-based approach by Miller *et al*^10^ (see Methods for description of Miller set). We compared the MutFam and Miller gene sets both by comparing individual genes and by mapping the genes to protein networks to identify common processes. This allowed us to identify those genes more likely to be implicated in cancer, as they mapped to common processes highlighted by two independent methods.

Finally, we use representative domain structures from the MutFams to identify 3D clusters of mutations for these families and show that this provides a strict method for filtering out passenger mutations leading to higher confidence driver gene set. These clusters are closer to both known functional sites and those predicted from highly conserved sites than non-clustered (i.e. unfiltered) mutations. A summary of the overall workflow is given in Figure 1.

**Figure 1.**
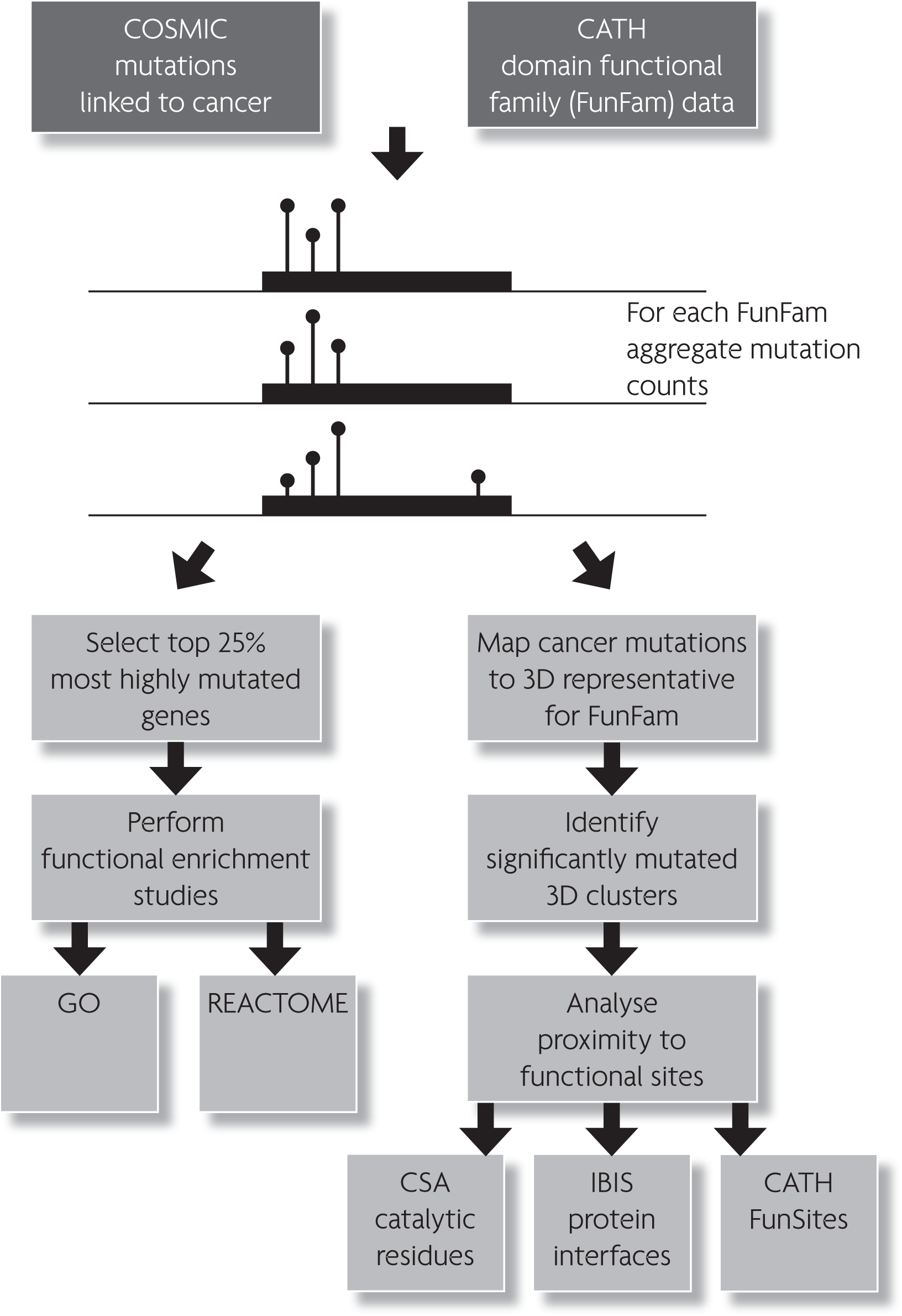
Flowchart summarises overall data processing and analysis pipeline.

### Identification of mutationally enriched domain functional families (MutFams)

To assess the value of MutFams for directly comparing diseases, and their functional signatures, we specifically analysed a set of 22 cancers identified in COSMIC. We grouped somatic missense mutations from 9,950 whole exome (or genome) samples in COSMIC v71 into 22 cancer types (based on tumour site, histology and respective subtypes) from which we identified a total of 259 MutFams (p < 0.05, permutation test with Benjami-Hochberg correction). For a summary of MutFams and number of samples by cancer type see **Supplementary Table 1**. The median number of MutFams per cancer is 13, ranging from 87 in Skin Cutaneous Melanoma to 2 in Uterine Carcinsarcoma. The number of MutFams per cancer does not correlate with sample size, i.e. the number of tumours sequenced (Pearson’s r = 0.232, p = 0.298; **Supplementary Figure 1**). We used neutral mutations (polymorphisms) from UniProt as a non-cancer control, resulting in 8 MutFams, with half of these belonging to Class II Major Histo-compatibility Antigens, for which a pool of variant alleles is likely to be advantageous^52^. **Supplementary Tables 2 & 3** summarise MutFams for 22 cancers and polymorphisms respectively, giving the UniProt functional keywords associated with each.

**Table 1.**
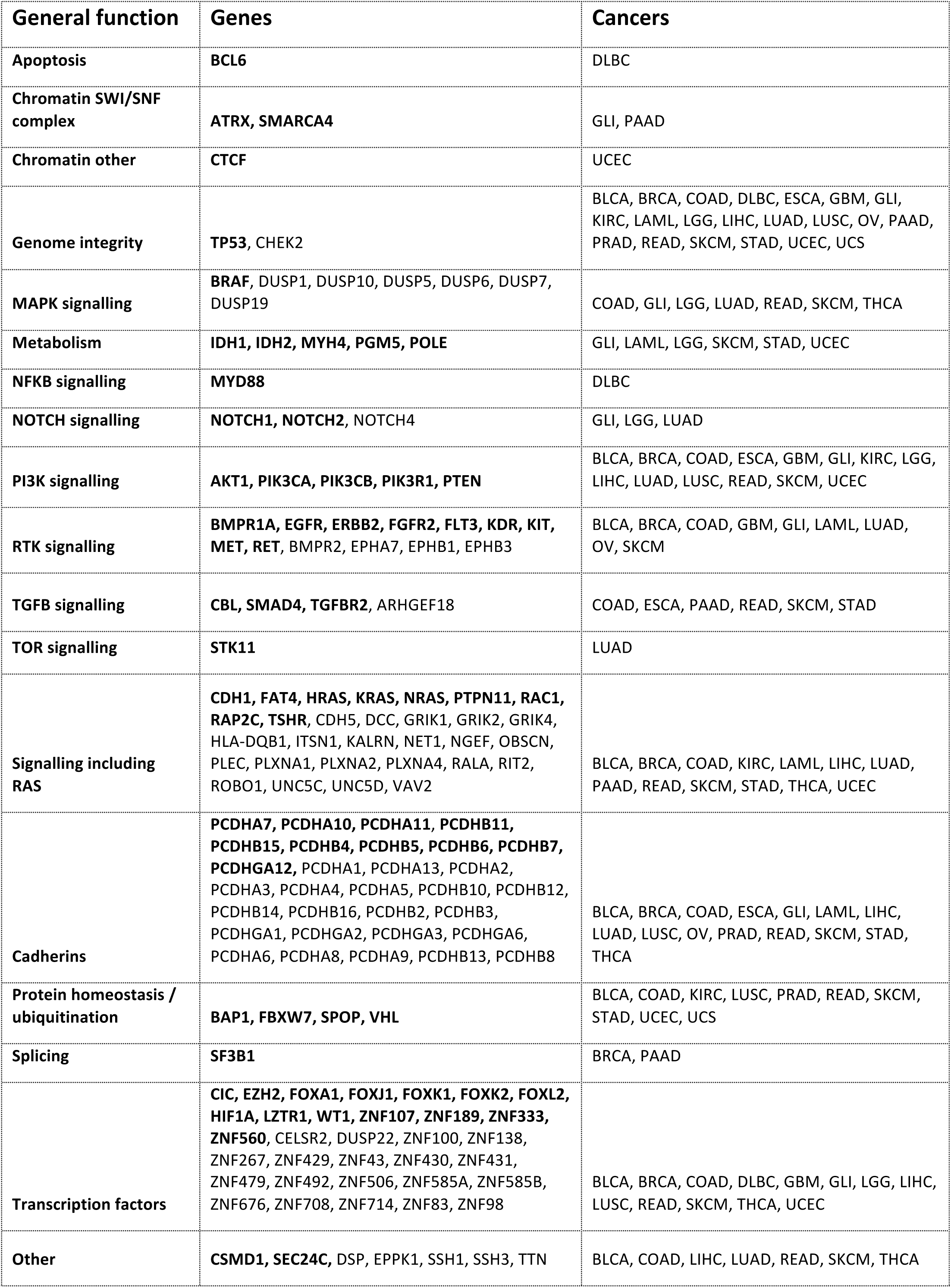
Summary of protein functions and cancers for the top mutated MutFam genes having some other supporting evidence. Genes in bold were also identified in either CGC or Miller.

We used hierarchical clustering of MutFams by cancer, based on shared MutFams, as shown in Figure 3. In agreement with other domain based methods^10,24,53,54^ mutations in the tumour suppressor p53 are commonly observed. We found 21 out of 22 cancer types have the MutFam ‘Cellular tumor antigen p53’. p53 mutations are found in many cancers^55,56^ but are not generally cancer-specific^57^. The second most common MutFam (11 out of 22 cancer types) is the tumour suppressor ‘phosphatase and tensin homolog’ (PTEN), a well-studied regulator of growth factor phospho-inositide signalling. Inhibition of PTEN has been shown to suppress regulation of the PIP3 secondary messenger, leading to increased growth factor signalling^58^.

**Figure 3.**
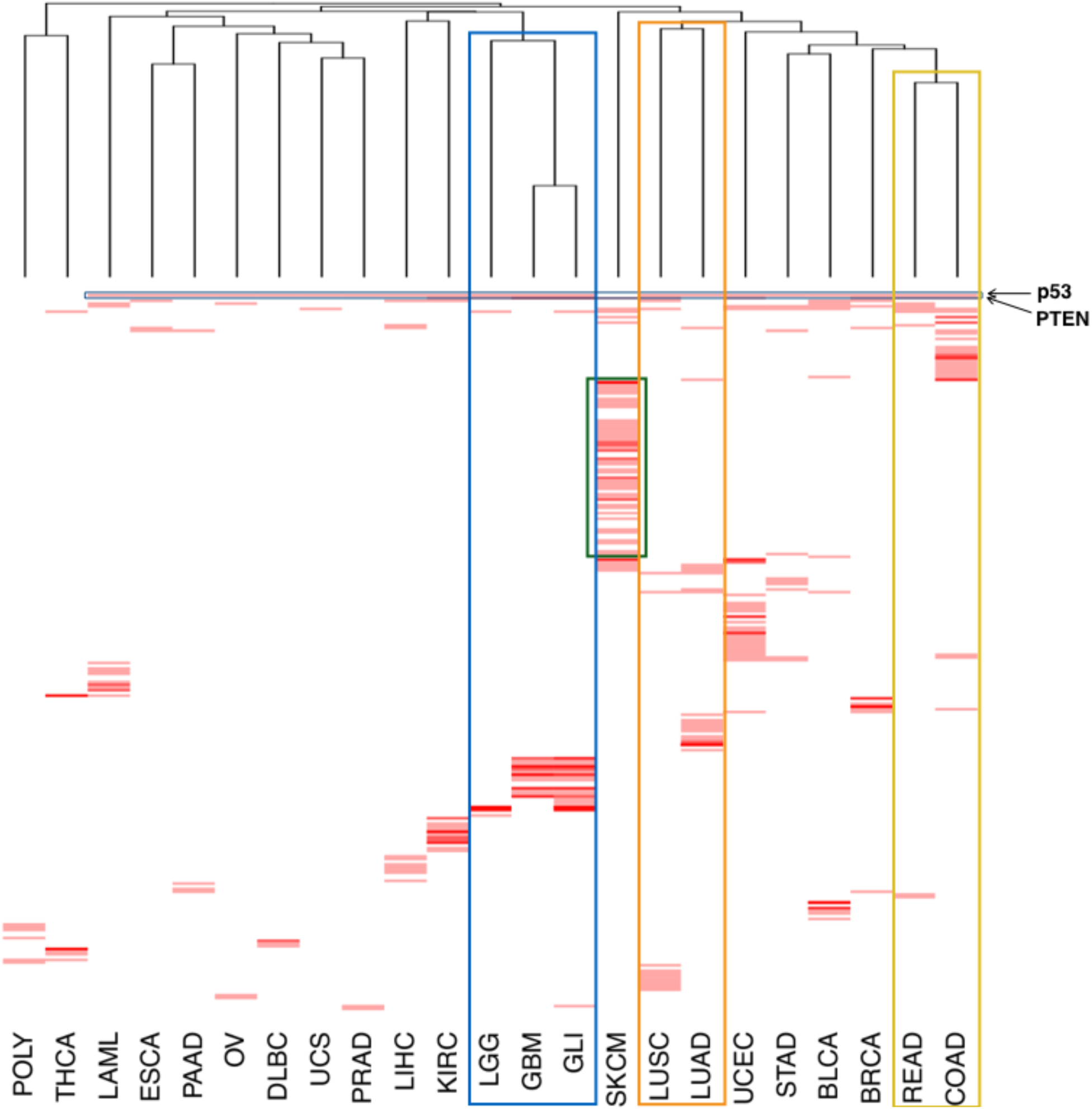
Heatmap of MutFam enrichments by cancer type. Each horizontal bar represents a MutFam, with higher enrichment factors shown in darker red (a full list of MutFams in given in Supplementary Table 2). Cancers found in the same site of the body and clustered by their MutFams are shown with rectangles: gliomas (blue), non-small cell lung cancers (orange) and colorectal adenocarcinomas (yellow). BLCA Bladder cancer; BRCA Breast invasive carcinoma; COAD Colon adenocarcinoma; DLBC Lymphoid Neoplasm Diffuse Large B-cell Lymphoma; ESCA Esophageal carcinoma; GBM Glioblastoma multiforme; GLI Gliomas; KIRC Kidney renal clear cell carcinoma; LAML Acute Myeloid Leukemia; LGG Low grade gliomas; LIHC Liver hepatocellular carcinoma; LUAD Lung adenocarcinoma; LUSC Lung squamous cell carcinoma; OV Ovarian serous cystadenocarcinoma; PAAD Pancreatic adenocarcinoma; PRAD Prostate adenocarcinoma; READ Rectum adenocarcinoma; SKCM Skin Cutaneous Melanoma; STAD Stomach adenocarcinoma; THCA Thyroid carcinoma; UCEC Uterine Corpus Endometrial Carcinoma; UCS Uterine Carcinosarcoma; POLY Polymorhisms (neutral mutations).

It can be seen in Figure 3 that cancers from the same site in the body (as defined by COSMIC) are found to cluster: those of brain tissues, low grade gliomas (LGG), glioblastoma multiforme (GBM) & gliomas (GLI); the two main histological subtypes of non-small cell lung cancer, lung adenocarcinoma (LUAD) and lung squamous cell carcinoma (LUSC) and the colorectal subtypes of colon adenocarcinoma (COAD) and rectal adenocarcinoma (READ). The large number of skin cutaneous melanoma (SKCM) MutFams (87) may be due to skin exposure to external carcinogens and is consistent with the highest mutation burden being observed in melanomas in general^59^. Additionally, thyroid carcinoma (THCA) is seen to cluster with the polymorphic dataset, being the only cancer not to have the p53 enriched MutFam. p53 is known to be affected in THCA via other mutation types (i.e. not missense) such as truncations, or by mutations occurring in regulatory transcription factors^60^. Additionally, while there are some MutFams common to all cancers, as discussed above, many are distinct (i.e. tissue-specific) and found uniquely within particular cancers, such as Skin Cutaneous Melanoma (SKCM), Acute Myeloid Leukemia (LAML), Bladder Cancer (BLCA), Liver hepatocellular carcinoma (LIHC) and Kidney renal clear cell carcinoma (KIRC) (as indicated in Figure 3; for specific MutFams see **Supplementary Table 2**).

#### MutFam genes are significantly enriched in CGC genes

By considering just Cancer Gene Census (CGC) genes that are observed to have missense mutations, we find that our list of MutFam genes is significantly enriched in CGC-curated cancer driver genes; the 49 drivers in common represent 21% of the MutFam genes identified (p < 10^−15^, Fisher’s). We compared this enrichment with that observed for the gene list identified by the related Pfam based approach by Miller *et al*^10^ (see Methods). The Miller set is also significantly enriched in CGC drivers with 30 genes in common representing 13% of the Miller dataset (*p*< 10^−15^, Fisher’s). Prior to any further filtering of these gene sets, each method identifies a majority of genes uniquely (Figure 4). We find 41 genes in common between the MutFam and Miller sets. Whilst this is a relatively low overlap between the genes identified by the MutFam and Miller sets, a recent study by Karchin *et al*^61^, which investigated overlap between driver genes predicted by 8 different methods, also showed that overlap of genes from individual methods with CGC was low. They did however find that the union of drivers from all methods showed significant enrichment of CGC genes. To investigate whether the unique genes in our dataset could be validated by other approaches, we examined their proximity in protein networks and also looked for association in biological pathways.

**Figure 4.**
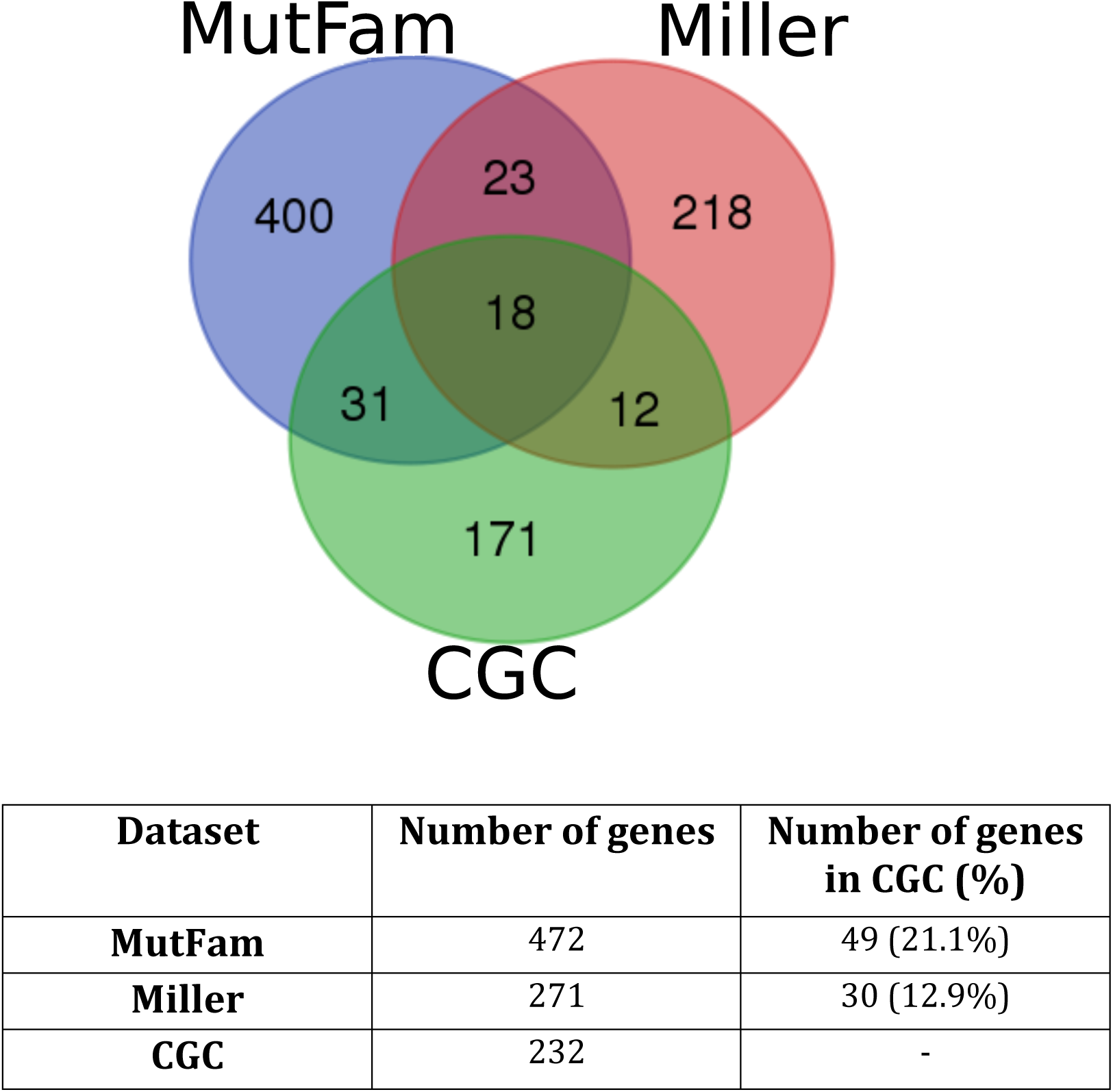
MutFam drivers are significantly enriched in CGC genes. The Pfam-based method (Miller) is also enriched in CGC genes and both methods predict distinct drivers.

#### MutFam genes are less dispersed than random on a protein-protein interaction network

Prior to any further filtering and enrichment of the top mutated genes from the MutFams, we analysed their overall dispersion on a comprehensive protein-protein interaction network using DS-Score, a measure adapted from Menche *et al*^42^ (see Methods). MutFam genes were significantly closer (less dispersed) on the network compared to randomly selected sets of genes (DS-Score 1.212, *p*< 1.2×10^−13^, Mann-Whitney test). Similarly, the set of genes from Miller also showed lower dispersion compared to random (DS-Score 1.463, *p*= 1.0×10^−8^, Mann-Whitney test) (**Supplementary Table 4**). These lower dispersion scores indicate that the tested genes are on average more closely connected in the network and thus more likely to be involved in the same biological processes.

### Enrichment of MutFam and Miller genes in common pathway modules and processes associated with cancer

We tested enrichment of the uniquely identified MutFam genes to identify whether they affected specific biological processes or were convergent on the same (or closely related) processes mediated by unique genes from the Miller set. We identified GO term enrichment using two approaches: (1) GO-Slim term enrichment on each of the gene sets and (2) GO Biological Process (GO-BP) term enrichment of network modules identified by mapping the gene sets to a pathway-based functional interaction network containing high quality curated annotations.

#### GO-Slim term enrichment

The 420 unique top mutated genes from the MutFams show enrichment in 16 GO-Slim terms, which we manually categorised into 5 cellular event categories: “embryonic development”, “cell migration”, “differentiation”, “stress response” and “cell signalling and transport” (Figure 5). The 224 Miller genes were enriched in 27 GO-Slim terms in 6 cellular event categories (**Supplementary Figure 2**). Despite differences between the specific GO-Slim terms found in each gene set, 3 of the cellular event categories were common to both MutFam and Miller (”embryonic development”, “cell migration” & “cell signalling and transport”) and each of these categories also contained at least one specific GO-Slim term common to both gene sets. The 12 common genes (between MutFam, Miller and CGC) were enriched in three GO-Slim terms involved in cell signalling and transport, including the immune related signalling term “I-KappaB Kinase/NF-KappaB cascade” and “receptor-mediated endocytosis” (**Supplementary Figure 2**). From this broad overview of the functional attributes of MutFam genes, we characterised specific processes in greater detail.

**Figure 2.**
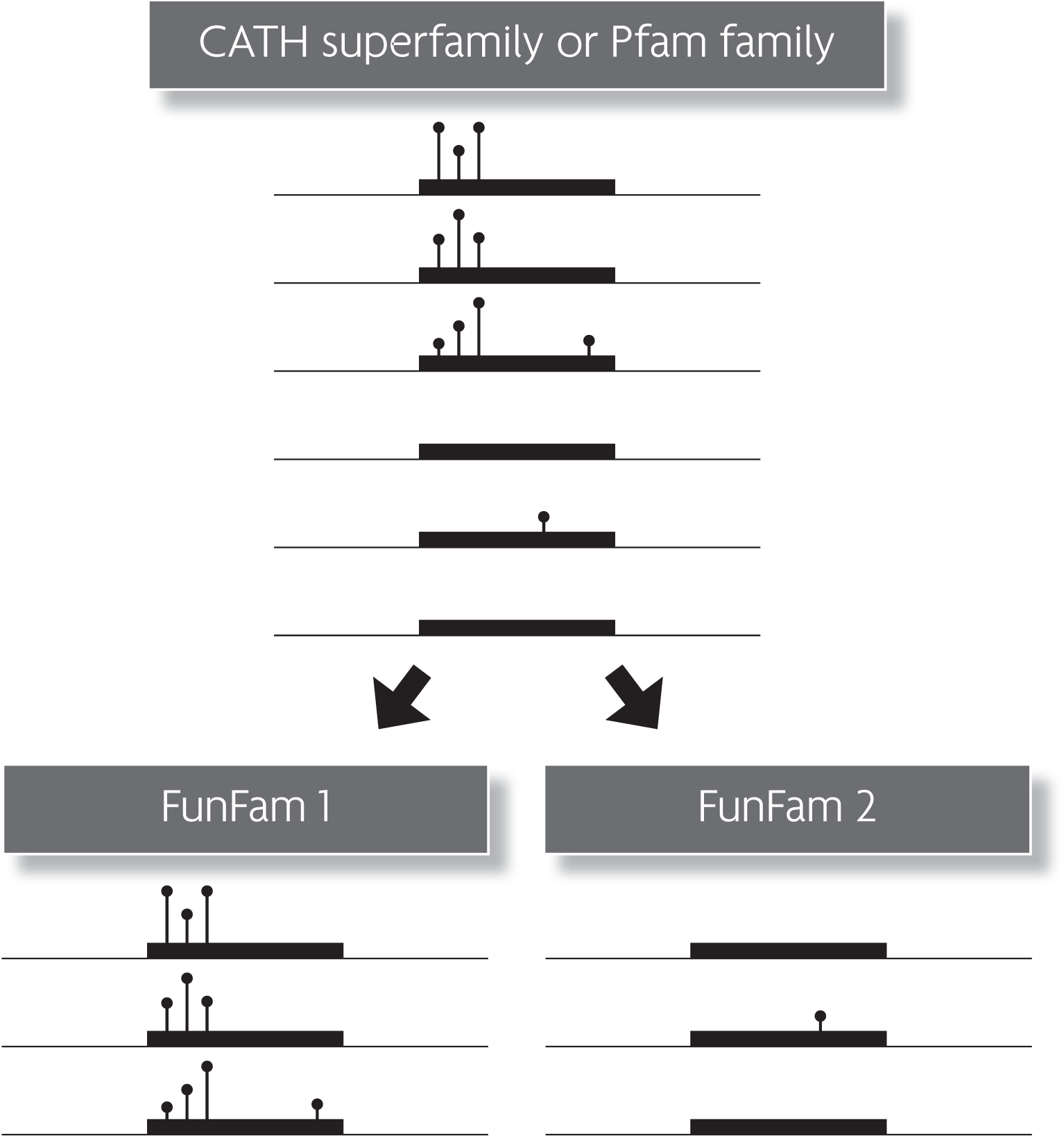
CATH-FunFams provide a more functionally specific classification than either Pfam family or CATH superfamilies. In FunFam 1, mutation enrichment is detected as significant but this might be diluted and missed if enrichment is calculated at the Pfam family level or the CATH superfamily level.

**Figure 5.**
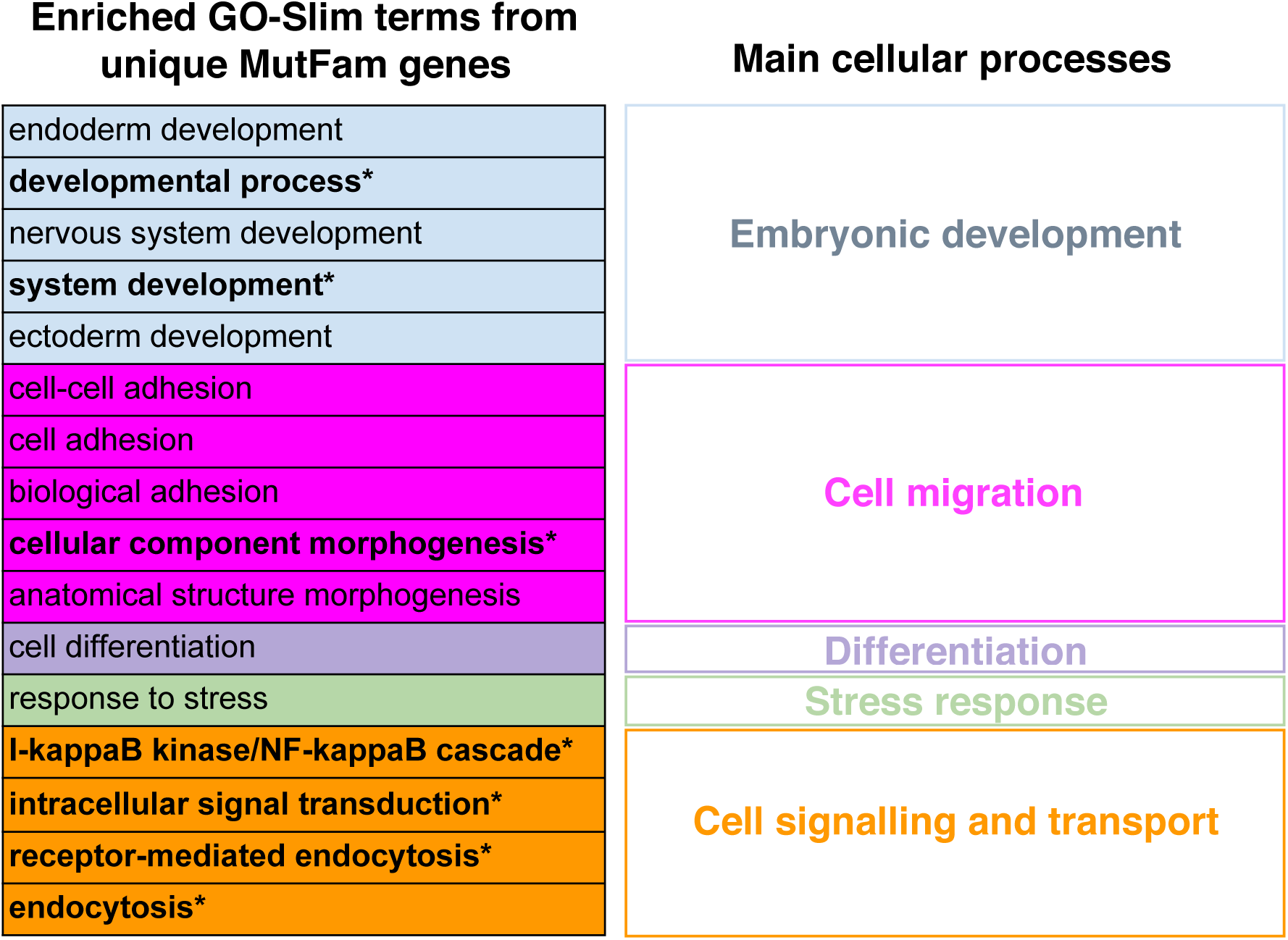
Genes uniquely identified from MutFams are enriched in 16 GO-Slim terms representing five main cellular processes. GO-Slim terms in common with Miller are shown in bold.

#### Analysis of the gene sets by functional network analysis to identify those more likely to be associated with cancer

Functionally related groups of genes were identified, for both the 12 common genes (”common”) and the genes unique to each of the datasets (”MutFam unique” etc.), by mapping each set to a pathway-based functional interaction network (Reactome FI, derived by Wu et al^62^), clustering these networks into modules and then calculating GO BP term enrichment for every module. Unlike the dispersion network measure outlined previously, which uses a protein-protein interaction network, Reactome FI uses the more specific pathway annotations of Reactome^63^ and allows better identification of functional relationships between proteins. We mapped 193 out of 420 (46%) unique MutFam genes, 119 out of 224 (53%) Miller genes, and 9 out of 12 (75%) common genes to the functional network. Not all genes map to functional modules because they lack any functional associations with other proteins in the network. Overall, 2 (out of 2) of the network modules for common genes, 12 (out of 21) modules for MutFams and 5 (out of 7) modules for Miller, showed enrichment for at least 1 GO BP term (FDR < 0.001).

The genes commonly identified by MutFam, Miller and CGC were enriched in receptor signalling processes and downstream kinase signalling pathways (see **Supplementary Section 1.1**). We found that genes uniquely identified by each method show different degrees of convergence at the functional module level. Such convergence may be very specific; a functional module identified from MutFam genes was associated with NOTCH signalling via NOTCH genes, whereas a module in Miller identified upstream and downstream regulators of the same pathway. Unique MutFam and Miller genes also converged on broader but clearly related biological processes of transcriptional regulation via proteins containing zinc finger domains. Finally, functional convergence was identified via broad and less specific pathways relating to cellular development (for further details on convergence between the unique MutFam and Miller genes see **Supplementary Section 1.1**).

Overall, our analysis of the genes identified by MutFam and Miller show that whilst there are only a modest number of genes commonly identified by both methods, at the level of biological processes and pathways there is much better agreement. Of the 193 MutFam genes mapped to the functional network, 168 were in enriched modules (˜85%). For at least 40% of these genes (68 genes) there was some other information supporting a cancer association i.e. literature, presence in a biological processes common with Miller genes or linked to cancer hallmarks **(See Supplementary Sections 1.1 - 1.3).** Thus by identifying different genes involved in common cancer-related pathways and by highlighting additional cancer-associated processes, the MutFam and Miller approaches can be considered broadly complementary.

Our pathway analysis further validates our MutFam selection, by revealing significant enrichment of MutFam genes in cancer-associated modules, and helps identify a more confident set of putative driver genes i.e. that are found both in mutationally enriched MutFams and enriched functional modules.

#### Detailed analysis of MutFam driver genes in brain cancers

In addition to the comprehensive analyses of MutFam mutated genes above, two specific cancers were subjected to more detailed analyses using Reactome Pathways and Gene Ontology (GO) enrichment: low-grade glioma (LGG) and glioblastoma multiforme (GBM). We found that processes enriched in GBM are consistent with it being a later stage (and more aggressive) glioma and include chaperone functions in proteostasis and immune functions. For a full summary see **Supplementary Section 1.4.**

### Identifying mutationally enriched 3D clusters in representative domain structures of MutFams

We have used functionally pure CATH-FunFams to detect domain families (MutFams) enriched in cancer disease mutations. To detect whether mutations within these MutFams were enriched in particular 3D locations we subsequently mapped the mutations onto a representative 3D structure for each family. Previous studies have shown considerable structural coherence within CATH-FunFams^43,50^. We used an in-house method (see Methods), to identify residues that have significantly more neighbouring mutations (i.e. within 5Å) than would be expected by chance. Therefore, by seeking for evidence of 3D clustering of mutations from MutFam genes we are able to identify a subset of putative driver genes and driver mutations with higher confidence to be associated with one or more cancers. We identified 175 clusters in 42 genes, comprising a total of 970 single point mutations. Since our hypothesis was that mutations clustering in 3D are more likely to be affecting functional sites in the protein, we assessed the value of these clusters for identifying putative drivers by their proximity to known and predicted functional sites in proteins. An example of a MutFam cluster occurring near functional sites in CHEK2 is shown in Figure 7.

**Figure 7.**
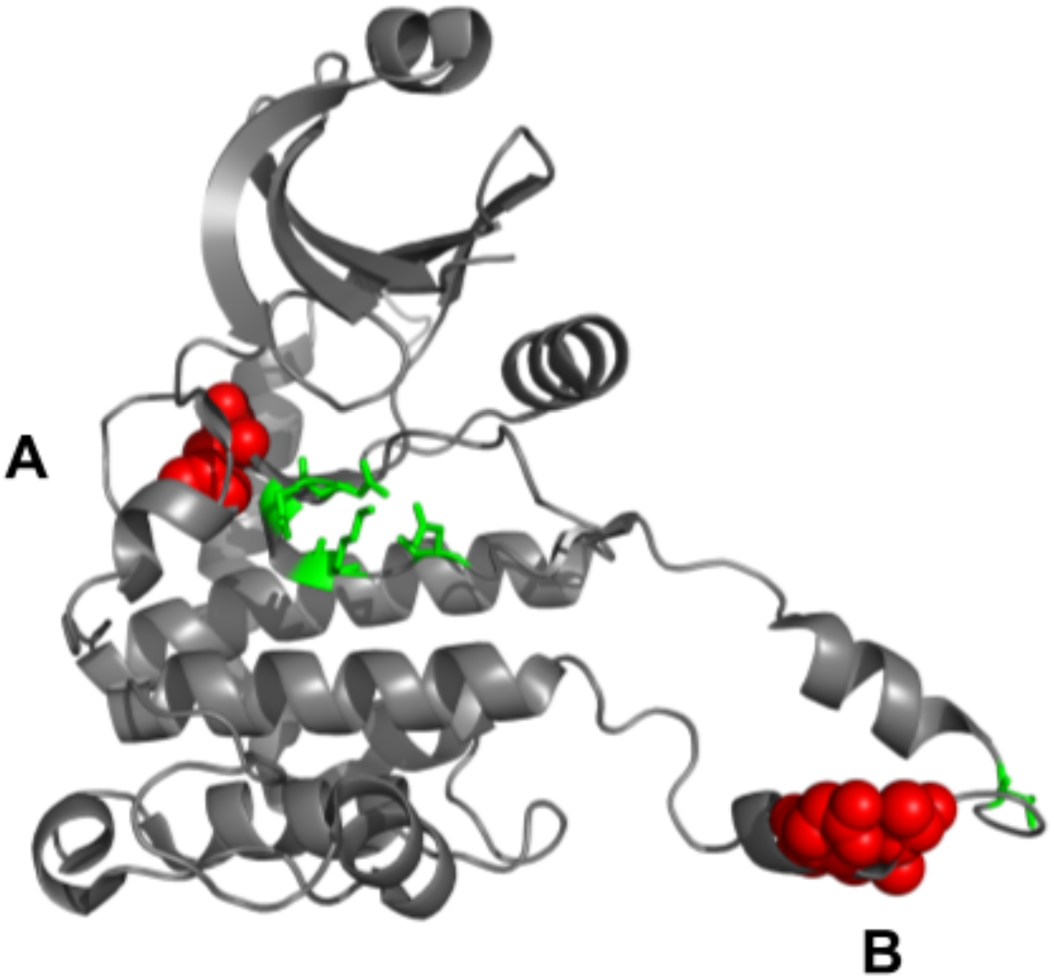
Mutations in CHEK2 cluster near specific functional sites. The phosphotransferase domain shown is representative of a MutFam identified in gliomas and glioblastoma “Calcium/calmodulin-dependent protein kinase type II”. Mutation clusters A and B (red) are within 5Å of both known and predicted functional sites (FunSites). Cluster A (centred on residue 355) is close to the ATP binding pocket, for which Catalytic Site Atlas (CSA) residues are highlighted green. Cluster B (centred around residues 392, 394 and 396) is in the activation loop near the APE motif, involved in kinase activation and function during which the loop moves towards the active site of the kinase; further CSA residues near cluster B are highlighted green.

#### Proximity of MutFam cluster mutations to functional sites

We calculated proximity distributions of mutations to catalytic residues, protein-protein interface sites (PPI), ligand-binding and FunSites (highly conserved residue sites in CATH-FunFams, see Methods). Distance distributions of mutations to each of these site types were calculated for the MutFam cluster mutations and for an unfiltered set of mutations from COSMIC (see Methods). Mutations in oncogenes and TSGs were examined separately to see if oncogene mutations were more likely to be near functional sites. As mentioned above, since the protocol used to find the enriched clusters is designed to filter out noise by excluding passenger mutations, we expected mutations in these clusters to be closer, in general, to functional sites than mutations in unfiltered datasets. All were compared to the control distribution derived from UniProt neutral mutations, which were further filtered by excluding heavily mutated proteins (>50 mutations) to avoid bias. Distances of mutations to functional sites are shown as empirical cumulative density functions for each functional site type.

##### Catalytic sites

The mutations in oncogenes from the unfiltered pan-cancer set tend to be closer to catalytic residues than neutral mutations, though not significantly (*OR* = 1.72,*p*<0.054, Fisher’s exact test), while TSGs show an equivalent but significant tendency (*OR* = 1.72,*p*<0.0044). However, MutFam clustered mutations do show a much more significant tendency to be closer to catalytic residues (*OR* = 2.98,*p*< 3.4×10^−15^) (see Figure 6).

**Figure 6.**
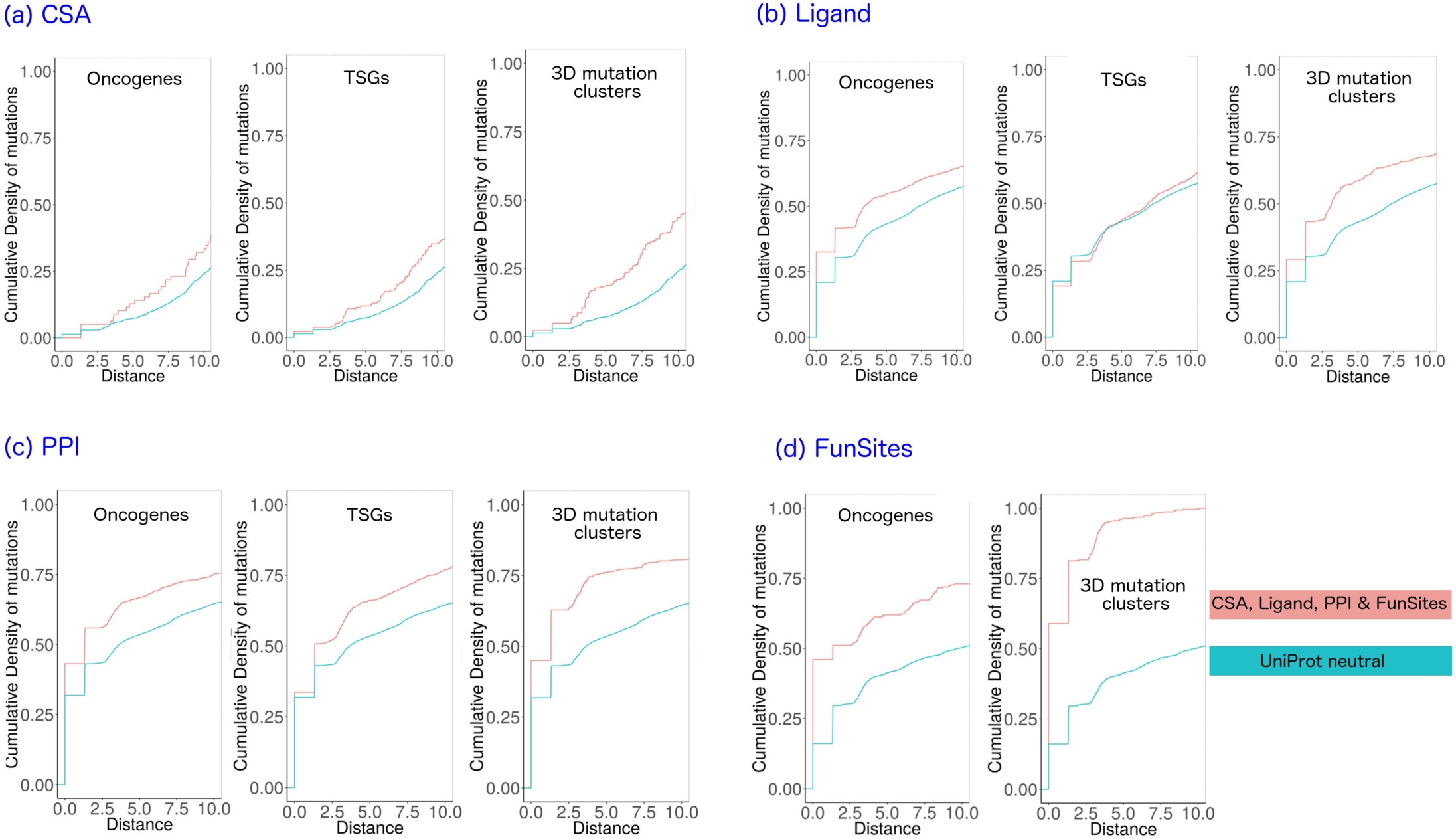
Proximity distributions show that 3D clustered cancer mutations are closer to functional sites than unfiltered mutations. For known functional sites (CSA, Ligand or PPI) 3D clustered mutations are closer than unfiltered mutations from either oncogenes or TSGs. For highly conserved residues in FunFam multiple sequence alignments highly enriched in known functional sites (FunSites), clustered mutations are significantly closer than unfiltered oncogene cancer mutations (there were too few distances measured to plot distributions for FunSites/TSGs). All distributions use UniProt neutral as a control.

##### Protein-protein interaction sites

Protein interfaces can cover large areas on the surface of the protein, thus substantial proportions of both cancer and neutral mutations are close to interfaces (Figure 6). Unfiltered cancer mutations in both oncogenes and TSGs show significantly higher propensities to lie close to interfaces (as defined by IBIS) than neutral mutations (oncogenes *OR* = 1.71,*p*< 3.6×10^−13^; TSGs *OR* = 1.71,*p*<4.7×10^−15^, Fisher’s exact test). As with catalytic sites, the tendency is more pronounced for the MutFam mutations filtered by 3D clustering *(OR = 2.52,p*< *2.3*×10^−16^).

##### Ligand-binding sites

Mutations in oncogenes show a small, significant enrichment near ligand binding sites (*OR* = 1.46, p<9.4×10^−8^, Fisher’s exact test), but those in TSGs are not significantly different to neutral (*OR* = 1.1 p< 0.23). However, MutFam clustered mutations are significantly closer to ligand-binding sites than neutral (*OR* = 1.78, p< *2.3*×10^−16^) (Figure 6).

##### Predicted functional sites (FunSites)

Predicted FunSites (highly conserved residues in multiple sequence alignments of FunFam relatives) have been shown to be enriched in both catalytic and protein interface residues^64^. Therefore, it is not surprising that we see similar trends to those observed for these types of residues (i.e. CSA and IBIS interface) (Figure 6). In addition FunSites are likely to include other sites important for the stability, folding and any allosteric mechanisms. Both oncogenes and our clustered MutFam mutations show a high propensity to lie close to these sites (oncogenes *OR* =2.45, p< 3.6×10^−11^; MutFam clusters *OR* =84.7,p< *2.3*×10^−16^, Fisher’s Exact test), with, again, the filtered MutFam mutations showing a more significant tendency. This is consistent with studies that show that somatic cancer disease mutations occur close to conserved sites^17,65^. This result reinforces the validity of our 3D clustering strategy to remove noise and identify driver mutations. Furthermore, using our functionally pure CATH-FunFams to detect structurally and functionally significant sites through conservation analyses helps to reveal the functional significance of these mutations.

### MutFam genes with the greatest functional impacts suggest putative drivers

A set of 472 genes was initially identified by selecting the top mutated genes from each of the CATH-FunFams enriched in mutations (see Methods). This aggregation of mutations in sets of domains having related functions provides some evidence for likely impact on protein function. To filter this set further and obtain a set of putative driver genes we used multiple lines of evidence by identifying: (1) genes in common with those predicted using the Miller method; (2) genes in common with CGC; (3) genes in cancer-related GO functional modules (as discussed above, with further details in Supplementary Sections 1.1 - 1.3 and Methods) and (4) genes from MutFams containing a 3D cluster near a functional site. By using these multiple sources of evidence for the functional impact of putative MutFam driver genes in cancer we can identify a set of 151 more confidently predicted driver genes that are predicted from MutFams and have at least one of the other four pieces of supporting evidence outlined above. For a full list of all 472 MutFam genes and the associated confidence scores, see **Supplementary Table 11**. A summary of the 151 MutFam driver genes is provided in Table 1, with each gene categorised by general protein function and the cancer types in which the gene was identified.

Since assessing the number of putative driver genes in MutFams that were listed in CGC (downloaded September 2016), additional cancer drivers genes have been added, increasing the overlap of MutFam and CGC by 11 genes (CNTNAP2, DCC, EPHA7, FKBP9, ISX, PIK3CB, PREX2, PTPRT, TNC, ZNF429, ZNF479) (CGC 5th Feb 2018 download), representing 25.9% of our MutFam genes overlapping with CGC genes with missense mutations. One of these was also identified in the Miller set (PIK3CB). Of the 11 genes, 3 (PIK3CB, PREX2 and PTPRT) are in the new “Tier 1” category (there are 2 tiers), defined by the CGC as those with the strongest evidence for a role in cancer. We had identified EPHA7 in our set of putative drivers filtered to include genes having both pathway enrichment and 3D clustering (see Supplementary Table 11). Although having only 2 mutations within the FunFam boundaries of the gene EPHA7 itself, the MutFam (Ephrin type-B receptor 2) was found to be enriched based on 12 mutations found in Ovarian cancer across 8 genes within the FunFam. Five of these genes only contributed a single mutation each to the MutFam. Calling EPHA7 a predicted driver gene on the basis of these numbers would be dubious, but by combining other types of evidence for functional effects the case is stronger. By using the total set of pan-cancer mutations, a total of 46 mutations in this FunFam gave a significant cluster in 3D that was near a predicted functional site. Additionally, GO process enrichment identified EPHA7 as part of enriched processes in cellular development. Other putative drivers highlighted in Table 1 but not in CGC or Miller are in the MutFam “Hepatocyte growth factor receptor”. The genes PLXNA1, PLXNA2 and PLXNA4 are enriched in cellular development - axon guidance processes and 68 pan-cancer mutations show a significant clustering of mutations, near a predicted functional site (i.e. based on the most highly conserved positions within the FunFam) and an IBIS PPI site. A recent study indicates role for PLXNA1^66^ in pancreatic cancer cell lines in enhancing tumour proliferation and invasion.

A full list of 472 genes in all identified MutFams is provided, along with the supporting evidence and overall score in **Supplementary Table 11.**

## Discussion

We have exploited a set of domain functional families (CATH-FunFams) classified within CATH domain structure superfamilies. Relatives in these families are clustered on the basis of structural and functional similarity. Structural coherence has been validated by structural superposition of relatives with known structures in the PDB, whilst functional purity is reflected in the performance of the families for providing functional annotations for uncharacterised relatives^67,43^ and in the fact that highly conserved residue sites within the family are enriched in known functional sites residues including catalytic and ligand binding residues^43^.

Previous studies have demonstrated the value of looking for mutationally enriched regions within proteins on the premise that mutations in these regions could convey functional effects driving cancer. Whilst most studies have exploited Pfam families, here, we examine the benefit of using families that have been explicitly clustered on the basis of functional similarity. We identify 259 mutationally enriched families (MutFams). The lack of any correlation between the sample size (number of tumours) and the number of MutFams identified indicates that comparisons between cancer types are not confounded by the different numbers of tumours analysed in each, and we would expect further samples to increase the enrichment factor of existing MutFams, and (to a lesser extent) highlight a few new MutFams that pass over the significance threshold.

We observe that diseases can be clustered in a sensible manner on the basis of their MutFams, with diseases affecting similar tissues (e.g. gliomas) clustering closely on the basis of shared MutFams. Despite differences in the driver genes identified by other independent methods (e.g. the Miller dataset) and CGC there is substantial convergence of MutFam driver genes at the level of GO biological processes affected, as also shown in Baudot *et al*^68^. Broadly, MutFams are complementary to Pfam-based methods.

The lower overall dispersion of MutFam genes on a PPI network compared to random implied that the genes were closer together on the network and therefore likely to be in related processes. This suggested that a more detailed analysis using a pathway-derived functional network would help in identifying enriched functional modules. Literature analysis of these enriched modules revealed biological processes known or highly likely to be involved in cancer and showed that the MutFam and Miller gene sets tend to converge on similar biological processes despite relatively low overlap of the actual gene sets. Furthermore, the cancer sets identified by the MutFam and Miller methods were found to be biased towards different tissues, and the genes therefore operating in different cellular contexts, which may partly explain this low overlap.

Finally, we also looked at the proximity of mutation clusters to functional sites, where available, to provide additional evidence for a gene’s functional relevance.

Our results suggest that our FunFam protocol is complementary to a Pfam based approach. Pfam families are larger and less specific than CATH Functional Families with a broader sampling of biological sequence space, and analyses have shown greater functional divergence within Pfam families than FunFams^43^. This divergence may lead to some drivers being missed as functionally diverse relatives in the family may have no or few mutations (as illustrated in Figure 2). By contrast the smaller size of FunFams may mean that enrichment values fall below the threshold, again causing driver genes to be missed. However, the higher number of genes identified by our MutFam analysis (i.e. 472 compared to 271 for the Miller set) suggests that the higher functional coherence of our families is helpful in detecting mutationally enriched families. For example, our MutFam approach detects EPHA7, which is not picked up in the Miller set, but was confirmed by a later release of the CGC. It is also reassuring that a high percentage of MutFam genes, mapped to the functional network, are in enriched modules. Furthermore, functional coherence of FunFams, and their associated structural coherence^43^, facilitates more accurate multiple sequence alignments which in turn facilitates more accurate prediction of functional sites, based on highly conserved positions within the FunFam. The functional site data, combined with available representative structures from FunFams, allow identification of mutations occurring (or clustering) near the known characterised functional sites or predicted functional sites. Since there are relatively few experimentally characterised functional sites, CATH-FunFams therefore bring significant benefits from the larger set of predicted functional sites they provide. Thus the use of representative structures from MutFams provides an effective filter for identifying mutations near functional sites that are likely to have specific functional effects.

In support of a combined MutFam and Pfam approach, Pfam includes domain families for which no structure is available and families for which there is no experimental characterisation of function (e.g. Domains of Unknown Function - DUFS), which are not currently represented in CATH-FunFams.

The complete list of 472 genes identified from MutFams is provided (**Supplementary Table 11**). By focussing on mutations that affect function i.e. by enrichment in CATH-FunFams, in biological processes linked to cancer, or in clusters near functional sites, we provide a novel protocol for filtering out passengers and predicting putative driver genes. In total, 151 genes are suggested by our MutFam method having biological process enrichment, 3D clustering or agreement with other datasets. A recent large-scale study by Bailey *et al*^69^ used a consensus scoring approach combining predictions from 26 software tools to identify 299 cancer driver genes. This set of 299 genes represents the most up-to-date consensus of drivers we had at the time of submitting this manuscript. Overall, a comparable 11% of 472 MutFam genes and 14% of 271 Miller genes are found in the consensus 299-gene set. However, 29% of the 151 MutFam gene set, i.e. those genes having evidence of functional effects, are in the consensus set, illustrating the value of our approach in filtering putative driver gene sets this way.

MutFams identified 79 genes uniquely that are not found in CGC or Miller, but have evidence for a functional role via biological process enrichment or 3D clustering. These could aid prioritisation of functional studies of putative novel driver genes identified from large-scale cancer genomics studies.

## Methods

### Datasets and sources

#### CATH Functional families

We used CATH-FunFams from version 4.0 of the CATH database^44^, which classifies 235,000 structural and over 25 million predicted domain sequences into 2,735 homologous superfamilies, which are then sub-divided into 110,439 functional families (CATH-FunFams).

CATH-FunFams are sets of evolutionary related domains clustered into families on the basis of predicted structural and functional similarity. CATH-FunFams are identified using agglomerative clustering of domain sequences within a CATH-Gene3D superfamily, and entropy based analyses to distinguish CATH-FunFams having distinct specificity determining residues^43^. CATH-Gene3D comprises all sequences from UniProt, which are known or predicted to belong to CATH domain structure superfamilies. CATH-FunFams have been highly ranked by the Critical Assessment of Functional Annotation (CAFA^67^) and shown to be more functionally coherent than Pfam families^43^. Typically several CATH-FunFams map to a Pfam family. In other words, although Pfam has been frequently used to increase the power of driver gene detection by accumulating mutation information across relatives within a Pfam family, this is also likely to introduce noise as Pfam families are not specifically classified for functional coherence and can contain relatives with rather diverse functions. Mutations in these domains may be exerting different effects as the genes may be operating in different pathway or cellular contexts and comprise different protein interfaces or active site residues. Previous studies comparing Pfam and CATH functional families for enzymes, showed that approximately 50% of Pfam families comprised more than one enzyme class (i.e. as reported by the enzyme classification) whilst less than 15% of CATH-FunFams comprised more than one enzyme class^43^.

For example, to illustrate the benefits of the CATH functional sub-classification, the schematic illustration in Figure 2 shows that analysis of the broad Pfam family shows no significant enrichment of mutations as some relatives have none or few mutations (for example, because these are paralogous relatives that operate in a different cellular context and hence lack the protein interface containing mutated residues found in the other relatives). Separating these relatives into distinct CATH-FunFams enables detection of mutational enrichment in CATH-FunFams. Conversely, the finer splitting of CATH superfamilies into FunFams may result in some CATH-FunFams being too small to detect mutation enrichment. It is therefore reasonable to suppose that mutation analysis by CATH-FunFams yields complementary driver gene lists to those obtained by related studies using Pfam families (e.g. by Miller *et al*, also analysed below).

Because of their functional purity, conserved residues within CATH-FunFams have been found to be enriched in known functional sites e.g. catalytic residues in CSA^26^. They have also been shown to be structurally coherent^64,70^. Highly conserved sites within CATH-FunFams (also described as FunSites) are found by analysing sequence conservation in a multiple sequence alignment of the FunFam, using the scorecons algorithm^71^. Scorecons conserved sites can only be reliably identified for CATH-FunFams having high information content (i.e. sequence diverse relatives) as measured by the DOPS score returned by scorecons. FunSite data was only generated for CATH-FunFams having a DOPS score ≥ 70 (range 0 to 100).

#### Cancer, polymorphism and disease datasets

*Cancer* datasets: 22 cancer-specific datasets were generated comprising somatic, non-synonymous missense exonic mutations from COSMIC^72^ v71, using variants from whole exome/genome studies, then filtering for each cancer type using tumour site and histology data with TCGA-style classes to define cancer types (summarised in **Supplementary Table 1**). Cancer datasets are used to define MutFams (see section “Calculation of MutFams - CATH-FunFams enriched in cancer mutations” below).

*UniProt neutral / polymorphism* dataset: 8,838 neutral mutations were obtained from 1,926 proteins using UniProt Humsavar^35^ (March 2014) by selecting entries annotated as “polymorphism”. This UniProt neutral dataset is used as a neutral control for the cancer datasets.

*Pan-cancer dataset:* 800,704 somatic, non-synonymous missense exonic variants from whole genome/exome studies from COSMIC v71 with no filtering by tumour site or histology. Note that the pan-cancer dataset is larger than all of the 22 cancer datasets combined as it includes many cancer sub-types from COSMIC that have few patient samples. This set is used to give as large and comprehensive cancer mutation dataset to use for 3D clustering as possible, based on the hypothesis that mutations from different cancers that cluster near the same functional site are likely to act via similar functional impacts. Subsets of the pan-cancer dataset were defined as “COSMIC oncogenes” and “COSMIC tumour suppressors” using gene roles identified by Wellcome-Sanger Cancer Gene Census (CGC)^51^. These sets allow for independent testing of the proximity of mutations to functional sites to account for any differences in the distribution of mutations found between oncogenes and TSGs.

#### PDB structures

Mutations were mapped to PDB structures using data in CATH v4.0 imported via SIFTS. Where multiple structures existed for a given UniProt protein, a single PDB was selected by selecting for maximum mapped sequence length followed by highest resolution, as per Stehr *et al* ^73^. Overall we were able to map 1,893 *COSMIC oncogene*, 3,184 *COSMIC TSG* and 8,838 *UniProt neutral* mutations to structures.

#### Functional sites

Functional sites were classified using catalytic residues from Catalytic Site Atlas 2.0 (CSA)^26^ and both protein-protein interaction (PPI) and ligand binding sites from NCBI-IBIS^74^. We additionally included ligand sites from ccPDB^75^ by filtering using their scoring metric (using AUC > 0.8 and MCC > 0.3) to include ligands ADP, ATP, FAD, FMN, GDP, HEM, NAD and PLP. We also analysed proximity to highly conserved sites (FunSites) within CATH-FunFams^43^. As previously described, FunSites are found using the scorecons algorithm^71^ for CATH-FunFams having high information content (i.e. sequence diverse relatives) as measured using the DOPS score returned by scorecons. FunSite data was only generated for CATH-FunFams having DOPS score ≥ 70 (range 0 to 100).

#### Calculation of MutFams - CATH-FunFams enriched in cancer mutations

For each of the cancer datasets (pan-cancer plus 22 cancer-specific) and UniProt neutral, we identified ‘MutFams’ as CATH FunFam domains significantly enriched in mutations. This protocol is based on Miller *et al* ^10^ and outlined below.

The enrichment of each FunFam domain is calculated as:

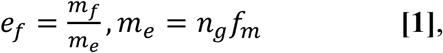

where *e*_*f*_ is the enrichment score of mutations (for a given database such as COSMIC) occurring in FunFam *f*; *m*_*f*_ is the observed number of mutations occurring in FunFam *f; m*_*e*_ is the expected number of mutations in FunFam *f*, calculated as total number of mutations observed in all genes containing the FunFam, *n*_*g*_, and the fraction of amino acids, *f*_*m*_, within CATH-FunFams compared to the total number in genes. Enrichment score significance was assessed using a permutation test as follows:

For each FunFam *f*:

Get the set of human genes containing the FunFam domain

> For each permutation *i* (1 to 10000):
>
> For each gene, *g*, count the number of mutations *n*_*g*_; randomly distribute the *n*_*g*_ mutations across gene *g*, allowing multiple mutations per amino acid.
>
> Calculate *m*_*i*,_ the total mutations for iteration *i* that are within the FunFam boundaries of the genes.

Enrichment *p*-value is defined as proportion of iterations where *m*_*i*_ *≥ m*_*f*_, where *m*_*f*_ is the observed mutation count for FunFam *f*.

For each disease type, MutFams were filtered to avoid noisy results from singleton mutations or very low counts by excluding those where total mutation count ≤ 10 (within FunFam boundaries, across all human genes and for multiple-spanning discontinuous sequence ranges, if applicable). Additionally, MutFams with enrichment factor *ef ≤ 1* were removed, as these are - by definition - not enriched. For example, out of the 3,124 CATH-FunFams in GBM (glioblastoma multiforme) with at least one mutation, 45 remain following the process and filtering outlined above. For each cancer type we applied Benjami-Hochberg (BH) correction to the permutation-derived *p*-values to correct for multiple testing of the mutation data set across multiple CATH-FunFams (applying FDR 5%). For GBM, 18 out of the 45 mutationally enriched CATH-FunFams remain following BH correction, leading to the set of 18 MutFams for GBM.

#### MutFam, Miller and CGC genes

*MutFam:* For each of the 22 cancers, we identified the top 25% of mutated genes for each MutFam. Our total MutFam gene set (n = 472) is the union of gene sets from all 22 cancers analysed. We then obtained sets of genes from other methods for comparison.

*CGC:* We used a subset of the full cancer driver gene list from Wellcome-Sanger Cancer Gene Census^51^ comprising those annotated as having missense mutations, resulting in 232 known driver genes; other mutation types may also have been annotated in these genes, e.g. frameshifts or deletions.

*Miller*: Genes identified from the Miller method^10^ obtained via download of the full results table from the MutationAligner website^20^. We then filtered the list to obtain those genes with significant domain hotspot residues (P < 0.05) resulting in 271 genes that contain at least one significant Pfam domain hotspot. We used this strategy because it gave a sensible number of genes that were deemed significant according to their analysis. The Miller set was chosen for comparison with MutFam as it is comprehensive and has been derived using a similar domain family based approach and because it provides the MutationAligner^76^ web resource for exploration and analysis of mutation hotspots in Pfam domain families. It has also been used in other comparative studies.

### Functional enrichment scores, network properties and protein structure analyses

#### Dispersion measure for genes mapped to a Protein-Protein Interaction (PPI) network

We measured the dispersion of genes on a network using DS-Score, adapted from Menche *et al*^42^ to give an overall proximity measurement for all of the genes in each dataset. Significance was assessed by comparison of the MutFam DS-Score with the average DS-Score obtained from 1000 sets of genes (n = 472, i.e. number of MutFam drivers) randomly selected from the STRING v10 protein-protein interaction network^41^.

#### GO term enrichment

Enrichment in GO-Slim terms was obtained for each gene set using the PANTHER online tool, testing for statistical over representation in GO-Slim Biological Process with P ≤ 0.01^77^.

#### Functional Interaction Networks

We analysed our MutFam gene set for pathway enrichment and pathway proximity. These MutFam gene sets were mapped to a Functional Interaction Network (FIN) using the ReactomeFIViz tool^62^, which provides a curated set of both known and predicted functional protein interactions. For each FIN, proteins were clustered into modules using the inbuilt community detection tool and each module analysed for enrichment in GO biological processes (FDR < 0.005). To compare the ability of different methods to identify putative driver genes, we compared the biological processes identified by using either MutFam or Miller gene sets. For an additional study focusing on gliomas, we obtained putative driver gene lists specific to the cancer types GLI, LGG and GBM and analysed these separately.

#### Calculation of mutations clustered on structures

We also separately calculated a set of pan-cancer MutFams using 800,704 somatic missense mutations from any whole exome or genome tumour sample, without any grouping by cancer type, to give 541 MutFams (*p < 0.05*, permutation test with Benjami-Hochberg correction). Note that this pan-cancer set is larger than the combination of the 22 cancer types, as it includes many rare cancers, each with very few tumour samples. We use this pan-cancer set to provide the largest possible dataset for 3D clustering, as mutations from different cancers clustering near a given functional site are likely to have similar functional impact. This larger dataset allows for increased detection of significantly enriched clusters. Mutations in the pan-cancer MutFams were mapped to their CATH representative domain PDB structures, where available. For the 541 MutFams in pan-cancer, 167 contained relatives with experimentally characterised structures. The representative structures selected from these FunFams had good resolution and the highest cumulative structural similarity to all other structures in the FunFam. For each representative structure, we tested for 3D clusters of mapped MutFam mutations. For each amino acid in the PDB chain with at least one mutation, we counted how many nearby (< 5Å) residues had at least one mutation, then tested whether nearby mutation counts for each residue were significantly different to the expected values arising from permutation testing. For each MutFam, we assigned all of the observed mutations to residues in the PDB structure at random and with equal probability, and then calculated the observed mutation counts near each residue as above. Following 5000 such permuted structure trials for each MutFam, 3D clusters were defined per MutFam as the residues having an observed nearby mutation count more than 2σ above the mean value for that residue over all permutations. Clustering was done using strict thresholds and only where at least one relative in the CATH-FunFam was structurally characterised. In future updates clustering would be performed using additional 3D models to expand the data set. We previously published this method (”MutClust”)^11^ and a similar method has also been independently described^78^.

#### Calculation of distance distributions of mutation and clusters to functional sites

We used an in-house method to measure the closest atomic distance between a mutated residue and a functional site residue; cumulative density function (CDF) distributions are plotted for these distances for each mutation dataset. Significance was tested with respect to neutral mutations for each disease type using Fisher’s exact test, with 8Å cut-off for mutation to site distance and *p < 0.01* considered significant (as used in Gao *et al*^36^). For clusters of mutations, proximity analysis was performed using the closest atomic distance between each residue defined as a mutation in a cluster and the nearest functional site residue. In addition to this filtering, for positions with multiple mutations, one mutation was selected at random as described in Gao *et al*^36^.

#### Odds ratio calculation

Odds ratios were calculated for the distance distributions of cancer mutations to functional sites by comparison with neutral mutations using Fisher’s exact test with 8Å as the upper bound for proximity; test based on Gao *et al*^79^.

### Scoring MutFam genes by predicted functional impact

The total MutFam gene set (472 genes) was annotated to find those most likely to have functional consequences using: (1) genes in common with those predicted using Miller method; (2) genes in common with CGC; (3) genes in cancer-related GO functional modules (defined below) and (4) genes from MutFams containing a 3D cluster near a functional site (defined above in section: “Calculation of mutations clustered on structures”). Additionally, as some MutFams occur in multiple cancer types and the MutFam enrichment factor is different in each, we calculated a mean enrichment factor for all cancers containing the given MutFam and used this to rank genes having the same score in the list of putative drivers.

*GO modules*: MutFam genes are classified as being in a GO module if present in any of the functionally enriched GO modules listed in **Supplementary Tables 5, 6, 7, 8, 12 or 13**. There are 71 genes classified as in MutFams and in a GO module.

## Acknowledgements

Paul Ashford was funded by BBSRC and Wellcome Trust. Camilla Pang was funded by Wellcome Trust. Tolulope Adeyelu was funded by the Government of Nigeria. Aurelio A. Moya-Garcí a received funding from the People Programme (Marie Curie Actions) of the European Union’s Seventh Framework Programme (FP7/2007–2013) under REA Grant Agreement No 623543.

## Author Contributions

P.A., C.S.M.P. and C.O. conceived and designed the study. P.A. and C.S.M.P. performed the experiments and analysed the results. T.A performed network dispersion analysis. P.A. and C.O. drafted the manuscript. A.M.G. provided constructive suggestions and discussions. All authors have read and agreed on the manuscript.

## Additional Information

Supplementary information accompanies this paper.

## Data Availability

All data generated during this study is included in this published article and its Supplementary Information files.

## Competing Interests

The authors declare no competing interests.

